# Changes in protein dynamical parameters derived from quasi-elastic neutron scattering spectra induced by coherent scattering: A molecular dynamics simulation study

**DOI:** 10.1101/2022.10.27.514007

**Authors:** Tatsuhito Matsuo

## Abstract

Protein dynamics at the sub-nanosecond timescale and the Å length-scale has widely been studied using quasi-elastic neutron scattering (QENS). In almost all QENS studies on hydrogenated proteins in D2O buffers, analysis of the spectra after buffer subtraction is carried out under the assumption that the remaining spectra arise from incoherent scattering of proteins while the contribution of coherent scattering is negligible. On the contrary, a study using polarization analysis has shown that the coherent scattering accounts for more than 10% of the total scattering intensity of hydrogenated proteins (Gaspar et al., Biochim. Biophys. Acta 1804:76–82 (2010)). In addition, the effects of coherent scattering on the values of dynamical parameters of proteins obtained by analysis of QENS spectra remain unclear. Here, molecular dynamics (MD) simulation on hen egg white lysozyme was used to investigate this issue. QENS spectra containing only incoherent scattering and those containing both incoherent and coherent scattering were calculated from the MD trajectory. Dynamical parameters were then extracted from the two simulated QENS spectra. Comparison of the resultant dynamical parameters has shown that the error in the values of the dynamical parameters induced by coherent scattering is at most 6%. This error is unlikely to significantly affect the results of QENS studies that investigate the relative changes in protein dynamics caused by different physicochemical conditions such as temperature unless dynamical parameters need to be determined with high precision at the absolute scale.

## Introduction

Quasi-elastic neutron scattering (QENS) is an experimental technique that has extensively been used to study dynamical behavior of proteins at the sub-nanosecond timescale and the Å length-scale [1–6]. The incoherent neutron scattering cross-section of a hydrogen (H) atom is more than 40 times larger than that of its isotope, deuterium, and any other atoms found in proteins [7], and about half of the total number of atoms constituting protein molecules are H atoms. As a result, experimentally measured QENS spectra are dominated by scattering of H atoms of the protein molecules. Because H atoms are distributed almost uniformly in space, dynamical information studied by QENS reflects the motions of H atoms averaged over all the H atoms in protein molecules.

In typical QENS studies of proteins, solution samples of hydrogenated proteins in D2O buffers are used as specimens. The QENS spectra are represented by the dynamic structure factor 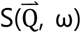, where 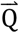 and ω are the momentum transfer and the energy transfer in the unit of ℏ of neutrons upon scattering, respectively. The spectra arising from protein molecules and their hydration water are obtained by subtracting the spectra of the buffer from those of the protein solution. The resultant spectra 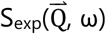 give almost flat 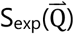, where 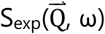 is integrated with regards to ω, and this is expected for incoherent scattering [8,9]. Since water molecules show a prominent broad peak arising from their coherent scattering at Q = 1.5–2.0 [Å^-1^] [8,9], the lack of such a peak in 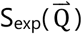 indicates that the contribution of coherent scattering is negligible in the system of hydrogenated proteins in D_2_O, and thus 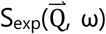 contains the dynamical information of protein molecules. Analysis of 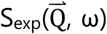 is, therefore, generally carried out with the assumption that it is equal to the incoherent dynamic structure factor 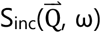 of protein molecules, for which various theoretical models and phenomenological fitting equations are formulated [1]. On the other hand, Gaspar et al. have experimentally shown by polarization analysis, which is a technique that can separate coherent and incoherent scattering [10], that there exists more than 10% of Q-dependent coherent contribution of the hydrogenated protein itself over a wide Q range in 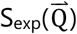 [11]. Although incoherent scattering is still the major component of 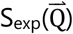, this seminal work has raised a concern about the validity of the dynamical information derived from the QENS study without polarization analysis. Therefore, it is of critical importance to investigate to what extent coherent scattering might affect dynamical parameters of protein molecules extracted from 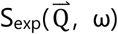 compared with those extracted from 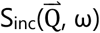.

In this study, this issue was investigated by simulating 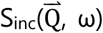 and 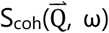 based on a MD trajectory of a hydrogenated hen egg white lysozyme (HEWL) in D_2_O. Dynamical parameters that are typically obtained and discussed in QENS studies were extracted from the simulated 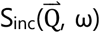 and 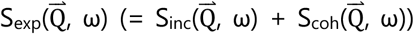. The resultant dynamical parameters were compared to reveal the possible changes in these values induced by coherent scattering.

## Materials and methods

### Molecular dynamics simulation

In this study, one of the trajectories of a hen egg white lysozyme (HEWL) molecule (PDB ID: 1AKI) generated previously was used [12]: The software GROMACS (version 2018.3) [13] was used to perform molecular dynamics (MD) simulation with the AMBER03 force field [14]. The heavy water model TIP3P-HW [15] was employed as a solvent molecule. In a cubic box, a HEWL molecule was placed and solvated such that all edges of the simulation box were ≥ 10 Å away from the protein surface, which simulates a protein molecule in solution. Counter ions were added to neutralize the charges of the entire system. van der Waals force and electrostatic interactions were cut off at 8.0 Å [14].

All the exchangeable (labile) hydrogen atoms of HEWL were replaced with deuterium by doubling the mass of the corresponding hydrogen atoms as in previous studies [16–18]. Whereas this simple procedure cannot explain an experimentally observed molecular behavior occurring at a long time scale (400 ns) [19], significant errors were not seen up to 50 ns from the start of the simulation. As a simulation time, 6 ns was chosen such that it was short enough not to generate large errors in protein dynamics and at the same time long enough to extract dynamical parameters obtained by QENS.

The system was first energy-minimized by the steepest descent method, followed by NVT and NPT equilibrations, which were carried out for 100 ps each with a time step of 2 fs. The resultant structure was subjected to a production run for 6 ns at 300 K and at 1 atm, and the first 5 ns of the trajectory was discarded to extract part of the trajectory which have reached a stable mean square displacement (MSD). Translational and rotational motions of the whole molecule were removed from the MD trajectory using the GROMACS command “gmx trjconv” so that only intramolecular motions are observed. The trajectory was saved every 2 ps.

### Calculation of incoherent and coherent dynamic structure factors from the MD trajectory

The incoherent dynamic structure factor 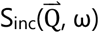 and the coherent dynamic structure factor 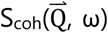 were calculated from the MD trajectory through the incoherent intermediate scattering function 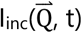 and the coherent intermediate scattering function 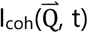, respectively, as follows:

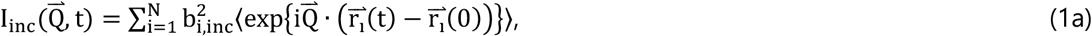

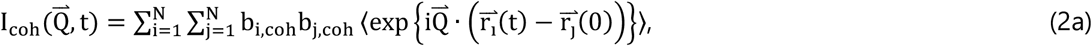

where N is the total number of atoms constituting a protein molecule, and b_i,inc_ and b_i,coh_ are the incoherent and coherent scattering lengths of the i-th atom, respectively, which are tabulated in Table 1. 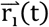 is the position vector of the i-th atom at time t, which is given by a MD trajectory, and 〈 〉 denotes the thermal average. In the case of solution samples, scattering intensity is orientationally averaged since protein molecules are randomly oriented, and the orientational average and the thermal average are independent. The vectors 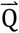 and 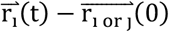 are thus treated as their modulus 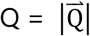 and 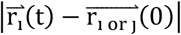, respectively. Eqs. 1a and 2a then read

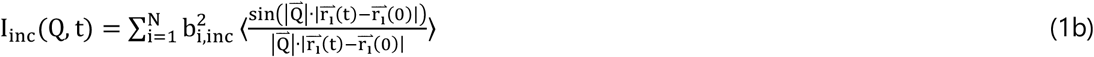

and

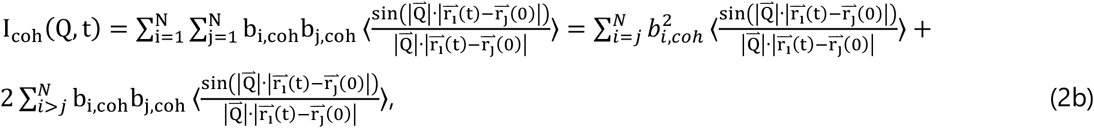

respectively.

**Table. 1.** Incoherent and coherent scattering lengths of atoms used in this study. The values are taken from the literature [1].

| Atom type | $b_{coh}$ [fm] | $b_{inc}$ [fm] |
| --- | --- | --- |
| H | -3.742 | 25.217 |
| D | 6.674 | 4.033 |
| C | 6.6484 | 0 |
| N | 9.36 | 1.98 |
| O | 5.805 | 0 |
| S | 2.847 | 0 |

S_inc_(Q, ω) and S_coh_(Q, ω) are the time Fourier transforms of I_inc_(Q,t) and I_Coh_(Q,t), respectively,

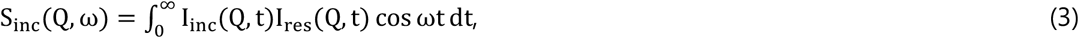

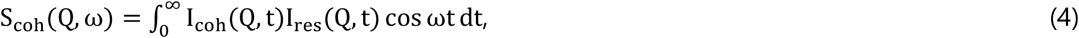

where I_res_(Q,t) is an instrumental resolution function, which defines the time-window of observation. In this study, this function was represented by a Gaussian function as,

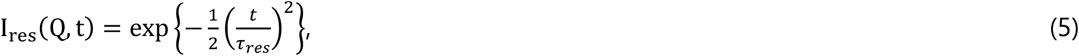

where *τ_res_* is the resolution time, which was set to 10 ps in this study. This corresponds to the energy resolution (FWHM) of 77 μeV. Since I_res_(Q,t) drops to almost zero at around t = 50 ps, I_inc_(Q,t) and I_Coh_(Q,t) were calculated up to t = 50 ps using the MD trajectory with the time length of 600 ps.

As stated above, S_exp_(Q, ω) is the sum of S_inc_(Q ω) and S_coh_(Q, ω) of protein molecules, i.e., S_exp_(Q, ω) = S_inc_(Q, ω) + S_coh_(Q, ω). Relative contributions of incoherent and coherent scattering to S_exp_(Q, ω) were investigated through the scattering intensities S_inc_(Q) and S_coh_(Q), where S_inc_(Q ω) and S_coh_(Q, ω) are integrated in the energy direction:

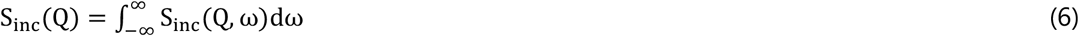

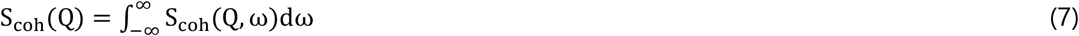

S_coh_(Q) corresponds to a scattering curve, which can also be obtained by small-angle neutron scattering. Because S_inc_(Q ω) and S_coh_(Q, ω) were simulated in the energy range between −0.5 meV and 0.5 meV, the integrations in Eqs. 6 and 7 were also conducted in this energy range.

### Analysis of the dynamic structure factors

Dynamical parameters such as the residence time and the jump diffusion coefficient, that are typically obtained by QENS measurements, were extracted by fitting the simulated S_exp_(Q, ω) and S_inc_(Q, ω) with the following phenomenological equation:

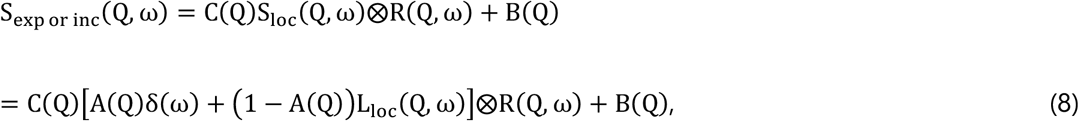

where C(Q) is a scaling factor between the fit and the simulated spectra, and S_loc_(Q, ω) represents the contribution from local atomic motions in a protein molecule. R(Q, ω) and B(Q) are a resolution function, which is a Fourier transform of Eq. 5, and a background, respectively. L_loc_(Q, ω) is a Lorentzian function describing local atomic motions, which is described as

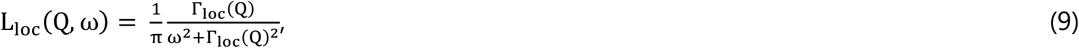

where Γ_loc_(Q) is the half width at the half maximum (HWHM) of the Lorentzian. The term including δ(ω) is the elastic component and that including L_loc_(Q,ω) is the quasi-elastic component.

Analysis of Q^2^-dependence of Γ_loc_ (Q) provides the detailed features of the corresponding diffusive motions. Γ_loc_(Q), which was obtained by the fitting of the simulated QENS spectra using Eq. 8, was found to be well described by the jump-diffusion model [20]:

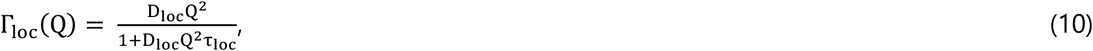

where τ_loc_ and D_loc_ are the residence time and the jump-diffusion coefficient of local atomic motions, respectively.

Q-dependence of A(Q), the elastic incoherent structure factor (EISF), provides information on the geometry of local atomic motions. The EISF was fitted by the following equation that describes a diffusion-inside-a-sphere model [21]:

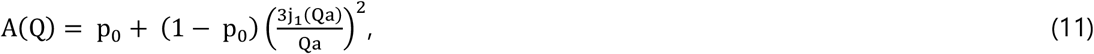

where p_0_ is the immobile fraction of atoms, motions of which are too slow to be resolved at the energy resolution employed, while (1 – p_0_) denotes a fraction of atoms resolved at the energy resolution employed. The latter atoms undergo diffusive motions within a sphere of a radius of a. j_1_(*x*) is the first-order spherical Bessel function of the first kind.

## Results and Discussion

First, S_inc_(Q) and S_coh_(Q) of HEWL calculated from the MD trajectory were compared to investigate the relative fractions of incoherent and coherent scattering in S_exp_(Q), which corresponds to the experimentally obtained S(Q) of proteins (S_exp_(Q) = S_inc_(Q) + S_coh_(Q)). The results are shown in Fig. 1 (a). It is seen that S_inc_(Q) is independent of Q values, which is consistent with experimental studies [8,9,11] while S_coh_(Q) shows a strong Q-dependence with small peaks at around Q = 0.6 and 1.4 Å^-1^. S_exp_(Q) thus shows slightly larger intensity below Q = 1.0 Å^-1^ than above Q = 1.0 Å^-1^, which is a tendency also observed in S_exp_(Q) obtained from the experimental QENS spectra on another protein troponin [9]. S_coh_(Q) corresponds to the so-called small-angle neutron scattering (SANS) curve, which is used for structural analysis of proteins, and hence it was compared with a SANS curve of a crystal structure of HEWL calculated using the program CRYSON [22] (Fig. 1 (b)). The resultant SANS curve agrees quite well with the simulated S_coh_(Q), suggesting the validity of the calculation procedure used in this study. Fig. 1 (c) shows the relative fractions of incoherent and coherent scattering in the simulated S_exp_(Q). It is found that the fraction of incoherent scattering depends on Q values, as reported previously using polarization analysis [11], and it reaches 80–90% as Q increases. Accordingly, the fraction of coherent scattering is 40% at Q = 0.4 [Å^-1^], above which analysis of QENS spectra is generally carried out, and then decreases to 10–20% at higher Q values.

**Figure 1.**
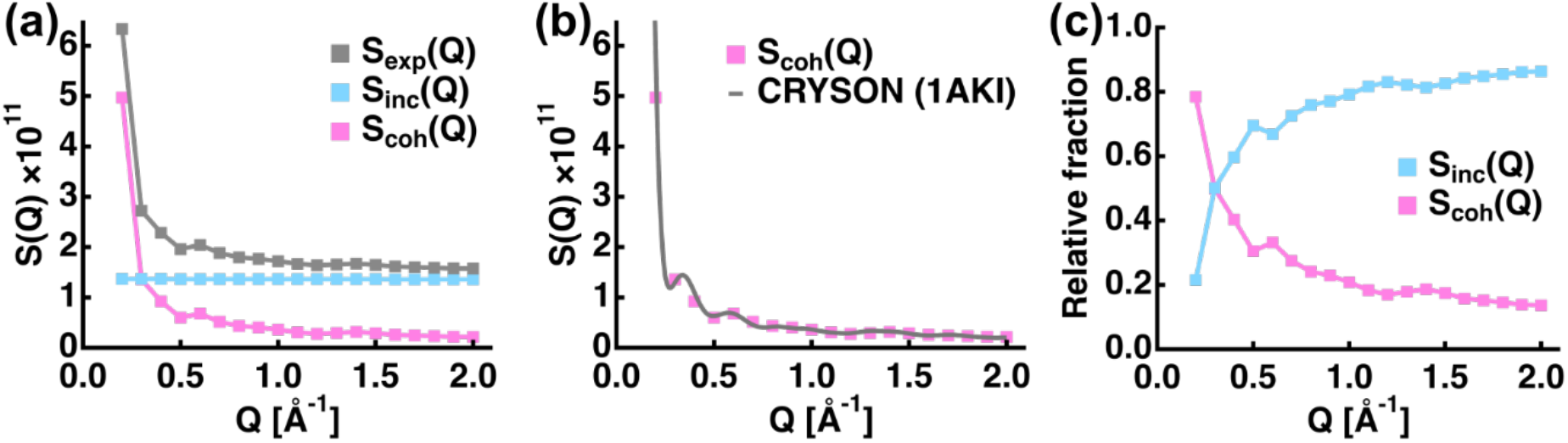
Q-dependencies of the S(Q) profiles of incoherent (S_inc_(Q)) and coherent (S_coh_(Q)) scattering calculated from the MD trajectory of hydrogenated hen egg white lysozyme (HEWL) in D_2_O. The profiles shown are from only HEWL molecules and do not contain contribution from hydration water and bulk water. (a): S_exp_(Q) denotes the sum of S_inc_(Q) and S_coh_(Q), which are shown in cyan and magenta, respectively. This is the S(Q) profile that is measured in typical QENS measurements without nuclear polarization. (b): Comparison of S_coh_(Q) calculated from the MD trajectory and that estimated from the software CRYSON, which calculates a small-angle neutron scattering curve, using a HEWL structure (PDB ID: 1AKI). (c) The relative fractions of coherent and incoherent scattering in S_exp_(Q) shown in (a) at each Q value.

Next, the shapes of the simulated S_inc_(Q, ω) and S_coh_(Q, ω) were directly compared, which has not previously been reported, neither experimentally nor theoretically. For this purpose, S_coh_(Q, ω) was normalized such that its peak intensity at ω = 0 [meV] is equal to that of S_inc_(Q, ω). The results are shown in Fig. 2 for Q = 0.5, 1.0, and 1.5 [Å^-1^]. It is seen that both spectra have essentially almost the same shape. On the other hand, when one takes a closer look at these spectra as shown in the insets of Fig. 2, the degree of broadening of the spectra is slightly different between S_inc_(Q, ω) and S_coh_(Q, ω) and this difference shows Q-dependence. To investigate the effects of these coherent scattering contributions on dynamical parameters derived from QENS measurements, analyses of the simulated S_exp_(Q, ω) and S_inc_(Q, ω) were carried out as a next step.

**Figure 2.**
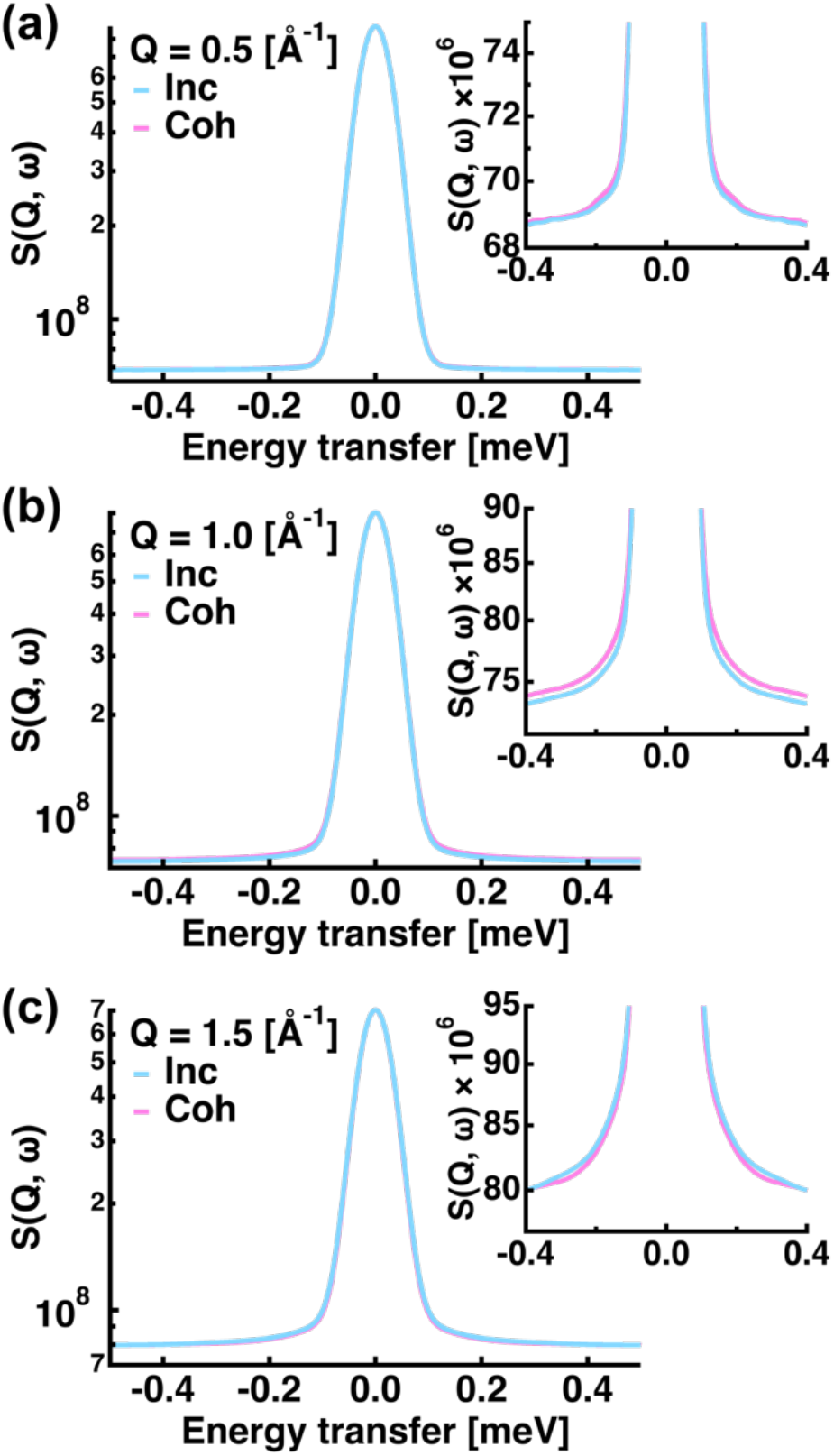
Comparisons of incoherent dynamic structure factor S_inc_(Q, ω) and coherent dynamic structure factor S_coh_(Q, ω) at Q = 0.5 [Å^-1^] (a), 1.0 [Å^-1^] (b), and 1.5 [Å^-1^] (c). Note that S_coh_(Q, ω) is normalized such that its peak is equal to that of S_inc_(Q, ω). The real fraction of Scoh(Q, ω) is as shown in Fig. 1 (a) and (b). The insets show the magnified views of the corresponding spectra.

The simulated S_exp_(Q, ω) and S_inc_(Q, ω) were fitted using the phenomenological equation of Eq. 8. A fitting example is shown in Fig. 3, which shows that both spectra are reproduced in good quality by this equation. The resultant Q^2^-dependences of the widths of the Lorentzian (Γ_loc_(Q)) and Q-dependences of the EISFs are summarized in Fig. 4. The Γ_loc_(Q) was found to be fitted by the jump-diffusion model (Eq. 10) in both cases as shown in Fig. 4 (a) and (c). Moreover, both EISFs were also reproduced by the diffusion-inside-a-sphere model (Eq. 11) as shown in Fig. 4 (b) and (d). Although S_exp_(Q, ω) contains S_coh_(Q, ω) in addition to S_inc_(Q, ω), slight differences in the broadening of the QENS spectra between S_inc_(Q, ω) and S_coh_(Q, ω), which was shown in Fig. 2, resulted in almost identical values of the Γ_loc_(Q) and the EISF as shown in Fig. 4 (e) and (f), respectively. Although the relative fraction of coherent scattering is 40% at Q = 0.4 [Å^-1^] (Fig. 1 (c)), this does not affect the Γ_loc_(Q) and the EISF values (Fig. 4 (e) and (f)). This is because the shape of the simulated QENS spectra is very similar between S_inc_(Q, ω) and S_coh_(Q, ω).

**Figure 3.**
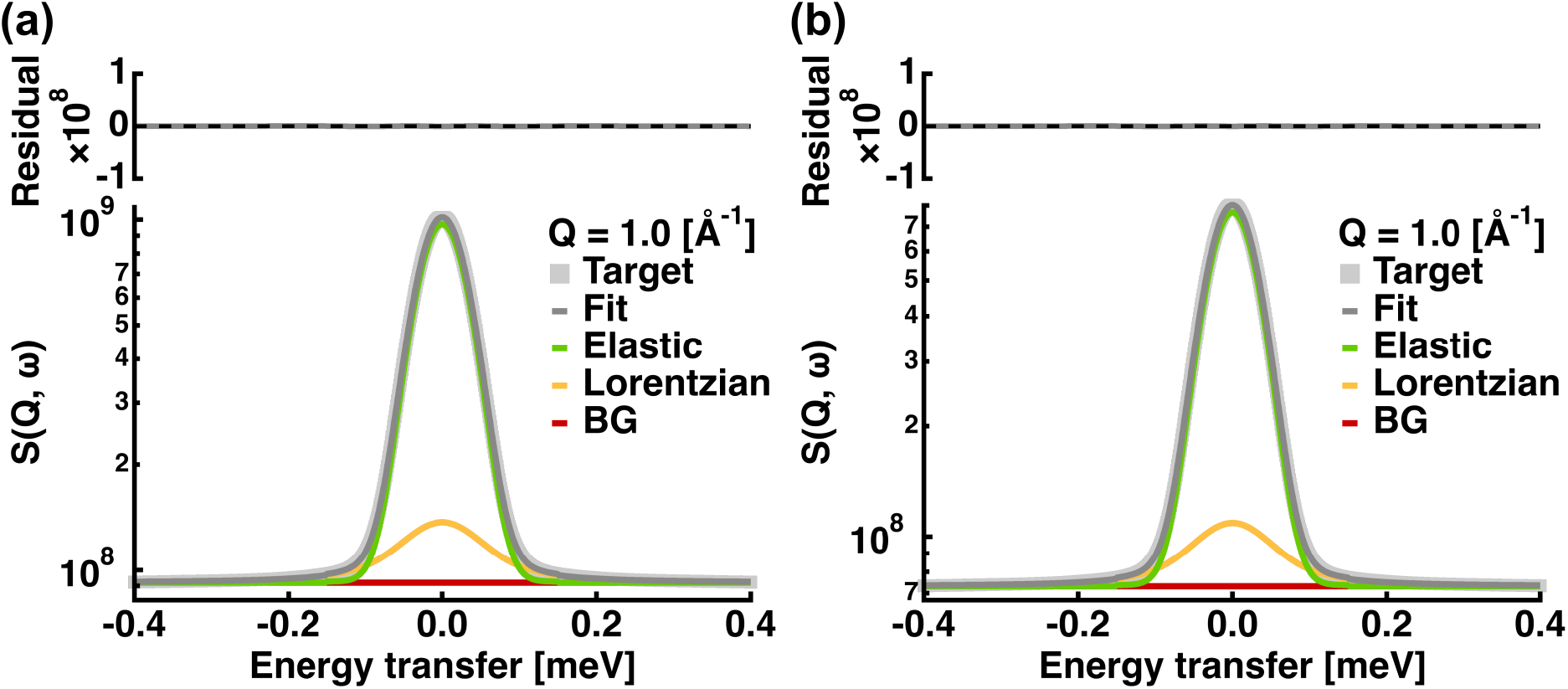
Examples of the fitting of simulated QENS spectra at Q = 1.0 [Å^-1^]. (a): QENS spectra containing both incoherent and coherent scattering, which is equal to those obtained from real measurements without nuclear polarization. (b): QENS spectra arising from only incoherent scattering. Grey filled squares represent the simulated QENS spectra, which are denoted as “Target”. The upper panels show the residuals between the “Target” and the fitted values.

**Figure 4.**
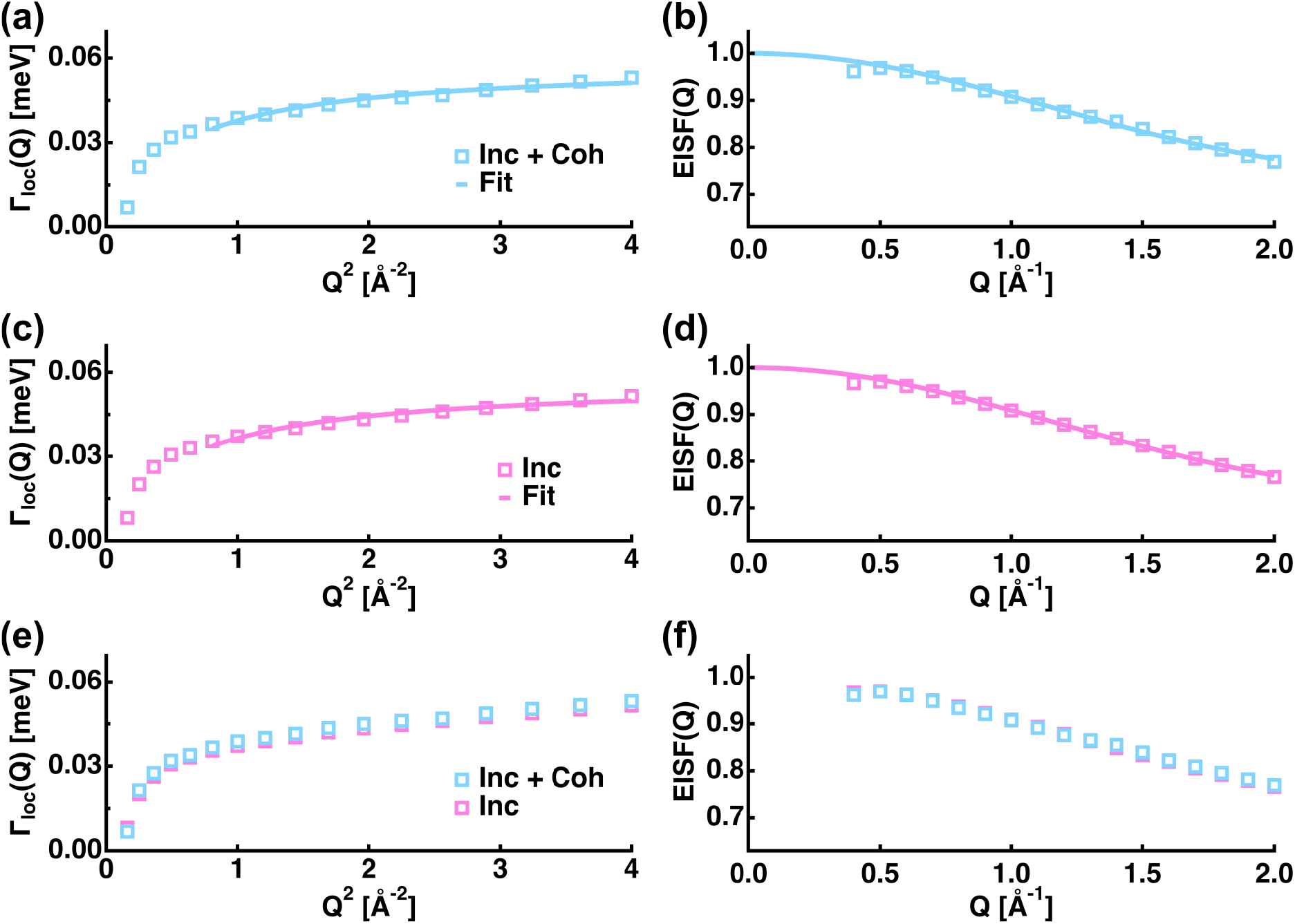
Q- or Q^2^-dependences of the widths (Γ_loc_(Q)) of the Lorentzian function the EISF. Open squares denote the Γ_loc_ values, which were estimated by the fitting of the simulated QENS spectra containing both incoherent and coherent scattering labeled as “Inc + Coh” (a, b) or those containing only incoherent scattering labeled as “Inc” (c, d). Solid lines denote the fits using Eq. 11 for the EISF and Eq. 10 for the widths. Comparisons of Γ_loc_ and the EISF between “Inc + Coh” and “Inc” are shown in (e) and (f), respectively.

Dynamical parameters obtained by the fitting of the simulated S_exp_(Q, ω) and S_inc_(Q, ω), i.e., the jump diffusion coefficient (D_loc_), the residence time (*τ*_loc_), the radius of a sphere (a), and the immobile fraction of atoms (p_0_), are compared in Fig. 5. It is found that the D_loc_ value increases from 1.54 ± 0.03 × 10^-5^ cm^2^/s to 1.63 ± 0.03 × 10^-5^ cm^2^/s by the contribution of coherent scattering, and *τ*_loc_ decreases from 11.67 ± 0.07 ps to 11.42 ± 0.07 ps. The a value increases from 1.389 ± 0.004 Å to 1.438 ± 0.003 Å while the p_0_ value increases from 0.722 ± 0.001 to 0.737 ± 0.001 by introducing coherent scattering. These results indicate that while the immobile fraction of atoms slightly increases, the contribution of coherent scattering to QENS spectra apparently leads to slightly enhanced protein dynamics.

**Figure 5.**
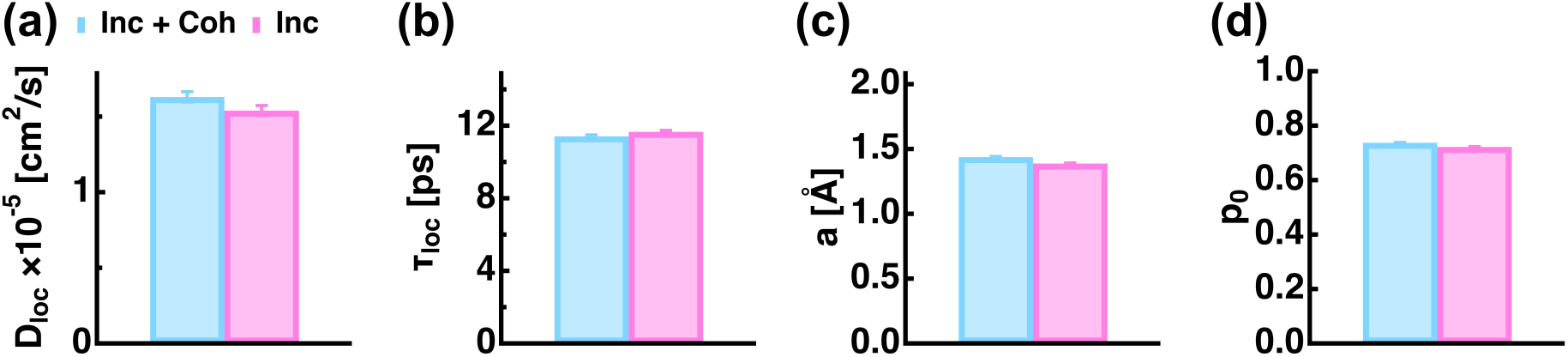
Summary of dynamical parameters extracted from the simulated QENS spectra. Comparisons of the jump diffusion coefficient (a), the residence time (b), the radius of a sphere (c), and the immobile fraction of atoms (d) are shown. Parameter values obtained from the simulated QENS spectra containing both incoherent and coherent scattering are shown in cyan while those obtained from the QENS spectra containing only incoherent scattering are shown in magenta.

Using modern neutron spectrometers such as BL02 DNA at J-PARC [23], dynamical parameters can now be estimated with a precision of 1–2% [9]. On the other hand, the differences in dynamical parameters caused by coherent scattering discussed in the previous paragraph are 5.8%, 2.2%, 3.5%, and 2.1% for D_loc_, *τ*_loc_, a, and p_0_, respectively. Thus, differences in some dynamical parameters induced by coherent scattering are beyond the precision of experiments and hence it is essential to bear in mind that dynamical parameters extracted from QENS spectra measured without polarization analysis might be varied by several percent compared with those that would be obtained from purely incoherent QENS spectra, which do not contain the contribution of coherent scattering. The present study thus suggests that in the case where dynamical parameters need to be determined at the absolute scale with high precision, QENS measurements with polarization analysis will be required as already proposed before [11]. On the other hand, in the case where changes in dynamical behavior of proteins induced by temperature [24,25], pressure [26,27], and other biological conditions [9,28], are investigated, the widths of the Lorentzian function and the EISF values generally show much larger changes than those shown in Fig. 4 (e) and (f) as the sample state is varied. In this case, a several percent of differences in dynamical parameters caused by coherent scattering would have negligible effects on the conclusion of QENS studies on proteins.

To the author’s knowledge, this is the first study to investigate the possible effects of coherent scattering on the dynamical parameters obtained by QENS spectra on proteins. In summary, analysis of the simulated QENS spectra, S_exp_(Q, ω) and S_inc_(Q, ω), has shown that despite the non-negligible contribution of coherent scattering, which accounts for 10–40% depending on the Q-range, the dynamical parameters extracted from the simulated S_inc_(Q, ω) are modulated by at most 6% by coherent scattering. However, these differences in dynamical parameter values are unlikely to affect the results of QENS studies to investigate the relative changes in protein dynamics caused by different physicochemical conditions of the sample unless dynamical parameters need to be determined with high precision at the absolute scale.

## Acknowledgements

The author is indebted to Prof. Judith Peters for reading the manuscript and fruitful comments.

## Notes

### Competing Interest Statement

The authors have declared no competing interest.

### Summary of Updates

Compared with the previous version, an additional paragraph was added to Introduction to better describe the background of this study.

